# Cryo-EM structure of RNA-induced tau fibrils reveals a small C-terminal core that may nucleate fibril formation

**DOI:** 10.1101/2022.01.28.478258

**Authors:** Romany Abskharon, Michael R. Sawaya, David R. Boyer, Qin Cao, Binh A. Nguyen, Duilio Cascio, David S. Eisenberg

## Abstract

In neurodegenerative diseases including Alzheimer’s and ALS, proteins that bind RNA are found in aggregated forms in autopsied brains. Evidence suggests that RNA aids nucleation of these pathological aggregates; however, the mechanism has not been investigated at the level of atomic structure. Here we present the 3.4 Å resolution structure of fibrils of full-length recombinant tau protein in the presence of RNA, determined by electron cryo-microscopy (cryoEM). The structure reveals the familiar in-register cross-β amyloid scaffold, but with a small fibril core spanning residues Glu391 to Ala426, a region disordered in the fuzzy coat in all previously studied tau polymorphs. RNA is bound on the fibril surface to the positively charged residues Arg406 and His407 and runs parallel to the fibril axis. The fibrils dissolve when RNAse is added, showing that RNA is necessary for fibril integrity. While this structure cannot exist simultaneously with the tau fibril structures extracted from patients’ brains, it could conceivably account for the nucleating effects of RNA cofactors followed by remodeling as fibrils mature.

**Significance statement:** Application of cryoEM has greatly expanded our understanding of atomic structures of mature pathological amyloid fibrils, but little is known at the molecular level of the initiation of fibril formation. RNA has been shown to be one cofactor for formation of fibrils of tau protein, and is known also to bind to other proteins, including TDP-43, FUS, and HNRNPA2, which form pathological inclusions. Our cryoEM structure of recombinant tau protein with RNA reveals a 36 residue, C-terminal fibril core bound to RNA which runs parallel to the fibril axis. We speculate that this structure could represent an early step in the formation of tau fibrils.

## Introduction

The two pathological hallmarks of Alzheimer’s disease (AD) are extracellular Aβ plaques and intracellular tau neurofibrillary tangles that accompany neuron loss in the brain (1). The deposition of tau aggregates in the brain is also the pathological hallmark of dozens of dementias and movement disorders known as tauopathies, including progressive supranuclear palsy (PSP), Pick’s Disease (PiD), chronic traumatic encephalopathy (CTE), and corticobasal degeneration (CBD) (2). In each of these conditions, monomeric tau proteins stack to form amyloid fibrils, but the trigger for fibrillization is poorly understood (3). Identical copies of tau stack into long β-sheets which in turn mate together tightly to form steric zippers; these fibrils are a common feature of all amyloid diseases (4).

Cofactors are necessary to form and stabilize tau fibrils *in vitro* and *in vivo*. *In vitro*, fibrillization of recombinant tau requires cofactors such as heparin, RNA, or arachidonic acid (5-7). Recent cryo-EM structures of heparin-induced tau filaments show they are heterogeneous and different from those of AD or PiD, which have larger fibril cores with different structures (8-10). *In vivo*, many cofactors are known to associate with NFTs in AD. These cofactors have been shown to stimulate AD-like phosphorylation of recombinant tau (5, 11, 12), and some of these cofactors are important for seeding activity *in vitro* (3, 5, 13). CryoEM structures of tau filaments extracted from the brains of patients with CTE, CBD, and PSP reveal residual density attributed to unknown cofactors: hydrophobic in CTE and anionic in CBD (14-16). These cofactors may stabilize particular tau fibril polymorphs (17) and together with post-translational modifications, govern which fibril polymorph dominates in the human brain (3, 18, 19).

RNA is implicated as a cofactor in the formation of fibrils of RNA binding proteins (RBPs) associated with amyotrophic lateral sclerosis (ALS), such as FUS, TDP-43, and hnRNPA1. These RBPs play important roles in gene expression by forming ribonucleoprotein complexes and participating in RNA processing steps including alternative splicing, stress granule formation and RNA degradation (20, 21). RBPs such as FUS and hnRNPA1 condense to form functional granules by liquid-liquid phase separation. Further aggregation, sometimes facilitated by RBP mutations, leads to pathological amyloid formation and accelerates development of neurodegenerative disease (2, 22-24). Notably, these proteins contain low complexity domains (LCDs) which further contribute to phase separation (22, 23, 25, 26) by a mechanism that is not yet clear.

RNA also induces tau to condense into a separate phase similar to RBPs (27) despite the fact that tau is not a *bona fide* RBP, nor does it contain an LCD. RNA forms a metastable complex with tau (18). RNA’s negatively charged phosphate backbone is thought to interact with the positive charge of the tau molecule to promote its aggregation (28, 29). These aggregates have pathological consequences. For example, RNA-containing tau aggregates in cell culture and mouse brains have been shown to alter pre-mRNA splicing (30). Moreover, crowding of tau molecules in the condensed phase facilitates tau stacking in a cross-β fashion to produce amyloid fibrils (6, 31). Similarly, RNA accelerates prion propagation *in vitro* and *in vivo* (32).

To learn how RNA interacts with tau at the atomic level and triggers fibril formation, we determined the cryo-EM structure of the fibril of recombinant full-length tau induced by RNA. With this structure and associated biochemical experiments we addressed several questions: (1) which tau residues comprise the RNA binding site? (2) why is the C-terminal region of tau poorly ordered in tau fibrils extracted from autopsied brains of patients with tauopathies? (3) how does RNA facilitate reversibility in fibril assembly and modulate fibril stability? (4) what role does RNA play in nucleating or deterring (33) formation of pathological tau fibril polymorphs?

## Results

### Generating full-length recombinant tau-fibrils

To test whether RNA can act as a cofactor to assist the fibril formation of full-length tau (residues 1 to 441, termed: Tau40), we incubated tau40 in the presence of total RNA extracted from mouse liver. We chose RNA from mouse liver because liver has less tau protein than other organs (34). In the presence of RNA, monomeric tau40 aggregates into amyloid fibrils (**Fig. 1A**). In contrast, no tau fibrils were observed in the absence of RNA, indicating that RNA induces these fibrils, and may be incorporated as a building block (**Fig. 1B**). To determine whether the fibrils contain full-length tau or a degraded product, possibly due to self-cleavage of tau, the fibrils were pelleted, washed and their molecular weight verified using SDS/PAGE. We observed that tau fibrils induced by RNA are primarily composed of full-length tau, with a small quantity of higher molecular weight species as shown in **Fig. 1C**.

**Fig. 1.**
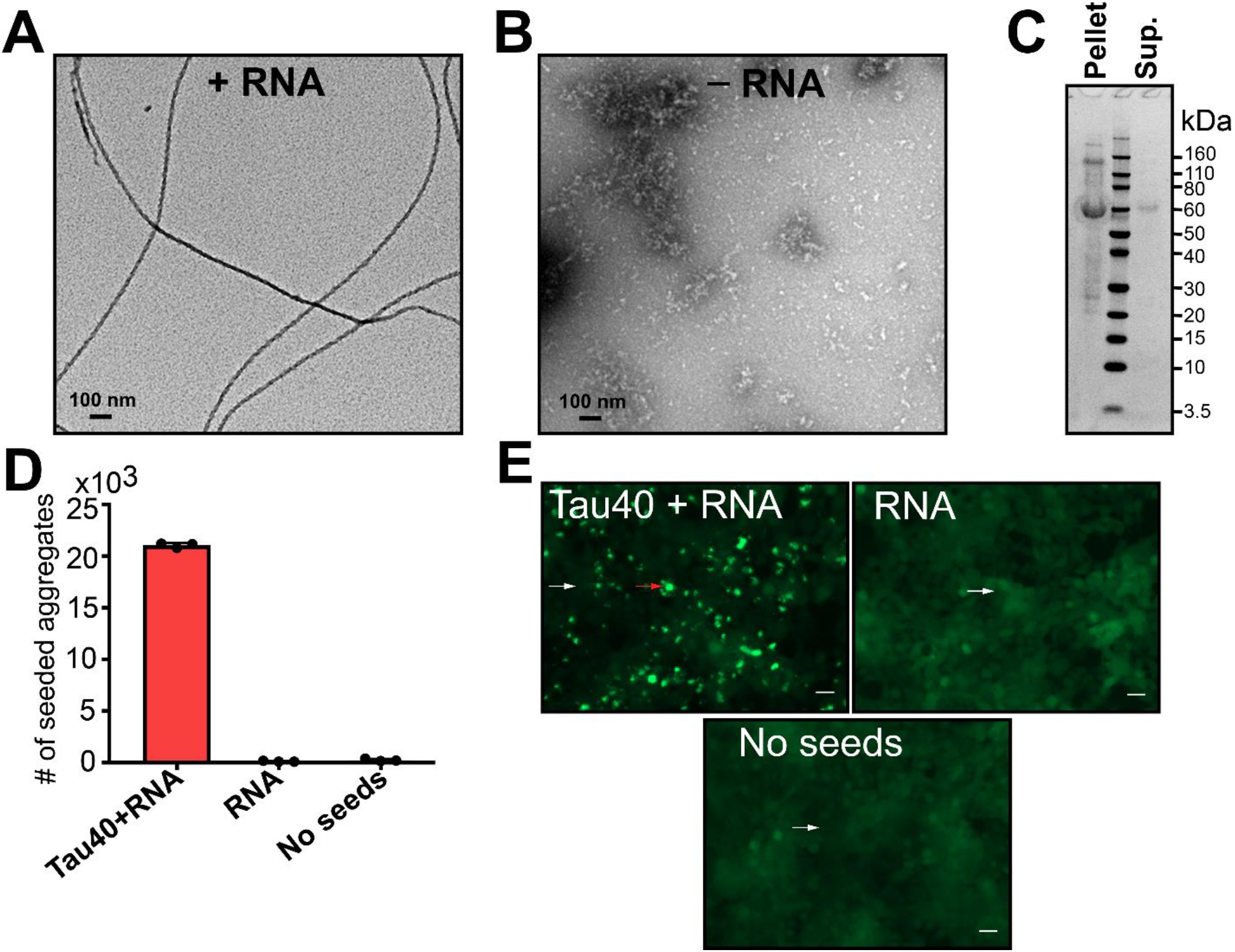
RNA induces formation of full-length tau amyloid fibrils. **(A)** Negative stain electron micrograph of tau40 fibrils grown in the presence of total RNA extracted from mouse liver. Tau fibrils were prepared by mixing 200 μl of 50 μM of monomeric tau40 in 20 mM ammonium acetate buffer pH7 with 400 μg/mL RNA and incubated with shaking for two days at 37 °C. **(B)** Electron micrograph of monomeric tau40 in the absence of RNA under the conditions of panel A. **(C)** SDS/PAGE analysis of tau-RNA fibrils after centrifugation at 159,000g and washing twice with RNA free water. **(D-E)** Quantification of the seeding activity of Tau-RNA fibrils, measured in HEK293 biosensor cells expressing YFP-tagged tau-K18. **(E)** Representative images of aggregates produced by seeding in HEK293 biosensor cells. Cells seeded with tau-RNA fibrils (left panel), and RNA only as a control (right panel). The red arrow highlights a cell representative of those that contain aggregates. The white arrows highlight cells representative of those that contain no aggregates. Scale bar 25 μm.

Next, we examined whether the tau-RNA fibrils have the capability to seed tau aggregation in HEK293 tau biosensor cells that express YFP-tagged tau-K18 (35). As shown in **Fig. 1D-E**, tau-RNA fibrils robustly seed tau aggregation in biosensor cells, whereas RNA alone displays essentially no seeding activity.

### RNA cofactor is essential for formation and stability of full-length tau fibrils

To test the role of RNA length on RNA-induced tau fibril formation, we performed *in vitro* fibrillation experiments using total RNA or RNA treated with RNAse (termed pre-digested RNA). Pre-digested RNA was incubated with tau40 monomers, but no amyloid fibrils formed even after 60 hours of shaking (**Fig. 2**). In contrast, undigested RNA induced spontaneous fibril formation over the same time course, suggesting the role of polymerized RNA as a cofactor in inducing tau aggregation.

**Fig. 2.**
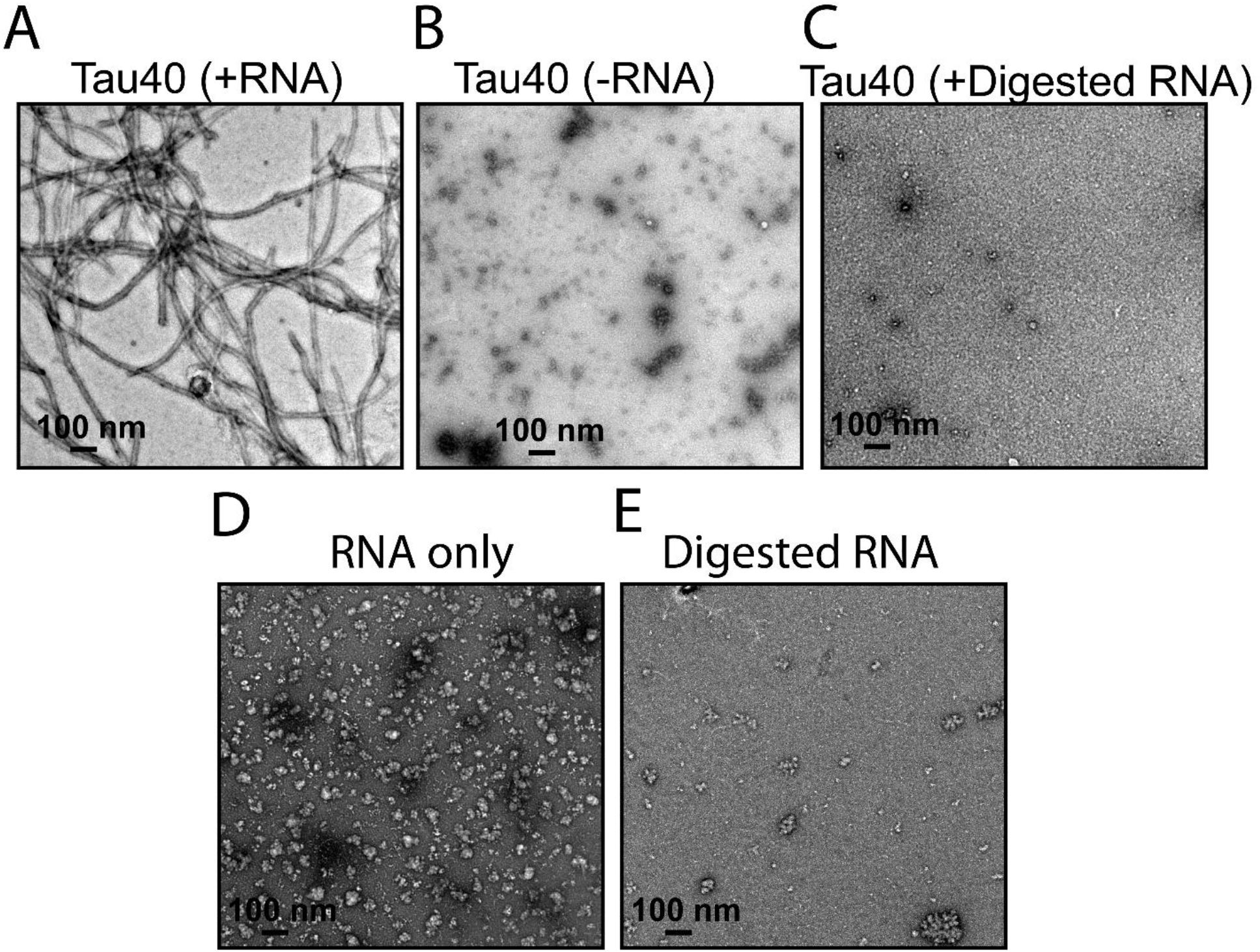
RNA enables *in vitro* fibril formation of tau. **(A)** Representative EM image of recombinant tau40 in the presence of RNA. **(B)** EM image of tau40 in the absence of RNA. **(C)** EM image of tau40 in the presence of pre-digested RNA. **(D)** Representative EM image of RNA only. (**E**) EM image of the digested RNA only. The digestion of RNA was performed by incubating the RNA with RNase at 4:1 ratio at 37°C for 6 hours. Scale bar 100 nm.

Fichou et al., reported that the presence of synthetic RNA (polyU) is required to sustain growth of fibrils of a truncated version of tau (termed: tau187, residues 255 to 441) even when the barrier to nucleation is removed by seeding with heparin-induced tau fibrils or mouse-derived fibrils (3). To investigate the influence of RNA on the seeding efficiency of tau, we used RNA-induced tau40 fibrils to seed monomeric tau in the presence of either pre-digested or undigested RNA, *in vitro*. We observed robust seeding of tau fibrils in the presence of undigested RNA sample (**Fig. S1 A**). Fewer and shorter fibrils are formed in the presence of digested RNA (**Fig. S1 B**). We didn’t observe any fibril in the presence of 5% seeds (2.5 μM of sonicated tau40 fibrils) and absence of RNA (**Fig. S1 C-D**), consistent with the critical role of polymerized RNA in supporting fibril formation. Similarly, no fibrils were observed when both RNA and seeds were predigested by RNAse (**Fig. S1 E**). We speculate that digested RNA would not have been sufficient to produce short tau fibrils if it were not for the small amount of RNA that accompanied the fibril seeds, consistent with the observation that seeding of monomer in the presence of digested RNA and digested seeds lacks even short fibrillar structures. These results suggest that polymerized RNA appears to be an essential constituent of self-assembling and seeding-active tau fibrils.

### RNA is the building blocks of tau fibrils

To assess biochemically whether RNA molecules remain associated with tau fibrils or dissociate upon fibril formation, we performed a co-sedimentation assay. We pelleted fibrils and washed them twice with RNA-free water. After sedimentation, pellets were resuspended in RNA-free water and RNA binding was evaluated by quantifying light absorption at 260 nm and performing RNA electrophoresis. These experiments reveal that RNA, predominantly of two molecular weighs, is incorporated into the fibril architecture (**Fig. S2**), consistent with previous studies that reported the binding of polyA RNA to tau-K18 and tau-K19 fibrils (29).

### Cryo-EM structure of full-length tau-RNA fibril bound to RNA

To discover how RNA induces tau aggregation, we determined the cryoEM structure of RNA-induced full-length tau fibrils to an overall resolution of 3.4 Å. We identified two fibril polymorphs in our cryoEM images and 2D classification: one fibril polymorph is twisted and the other lacks a twist. The twisted species was the more abundant, accounting for 75% of the fibrils. The species lacking a twist accounted for the remaining 25% of fibrils and was not suitable for structure determination. The twisted tau-RNA fibril exhibits a pitch of 829 Å. It is composed of two protofilaments related by a pseudo 2_1_ screw axis. It exhibits a helical twist of 179.16° and a helical rise of 2.4 Å (**Table 1**, **Fig. 3B and Fig. S3)**. The protofilaments are composed of flattened tau molecules stacked in parallel, in-register β-sheet alignment (**Fig. S4**). The fibril core spans 36 residues near the C-terminus of tau, from Glu391 to Ala426 and encompasses five β-strands (β1: Glu391-Val393, β2: Ser396-Ser400, β3: Asp402-Pro405, β4: His407-Ser413 and β5: Asp418-Val420) **(Fig. 3A & E)**. Notably, this region of tau has been characterized as a disordered fuzzy coat in all previously determined structures of tau fibrils, which includes the *ex-vivo* fibrils from AD, CBD, PSP, GGT and CTE patients as well as heparin-induced fibrils (8-10, 16, 36).

**Table 1:**
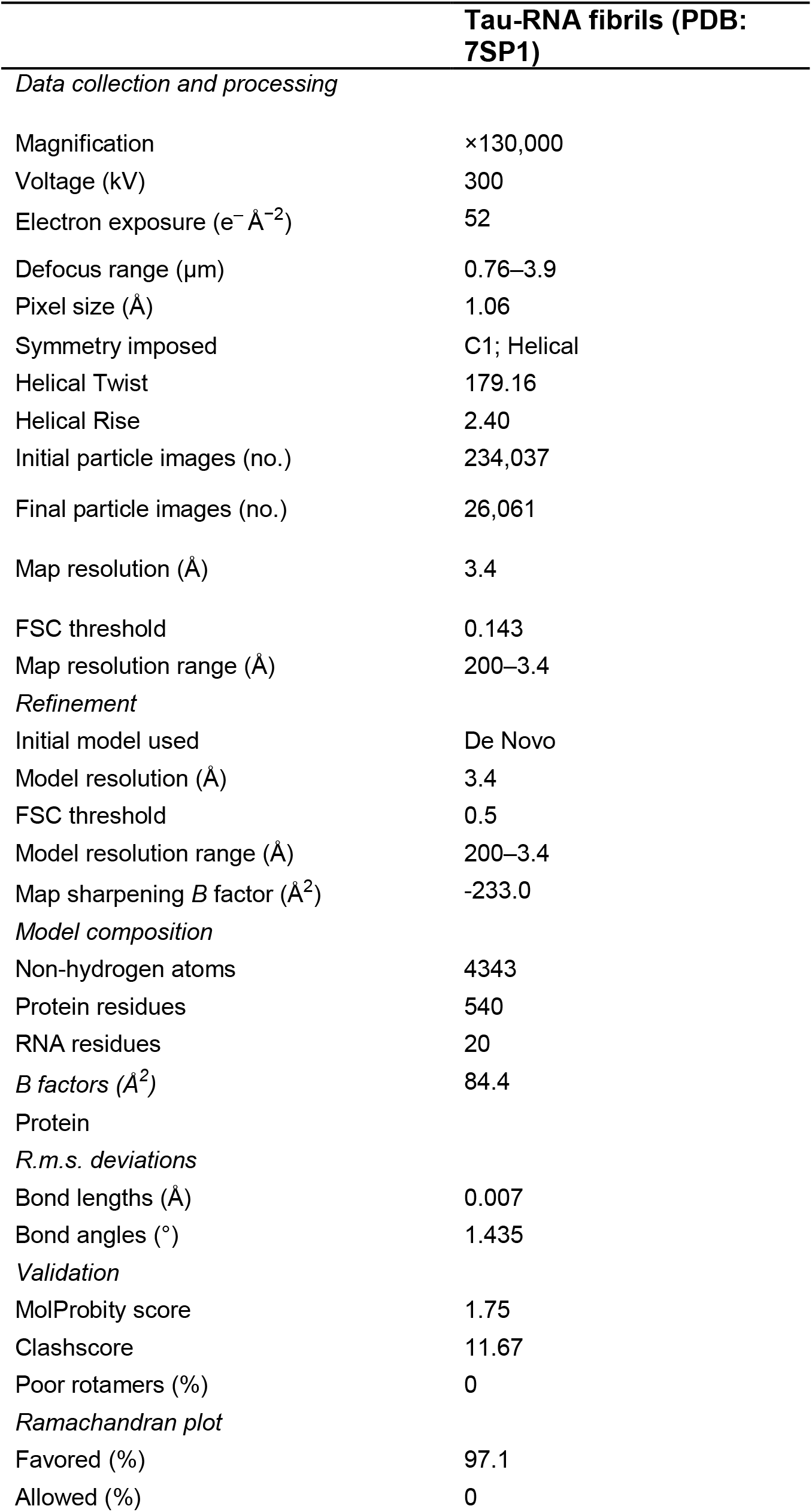
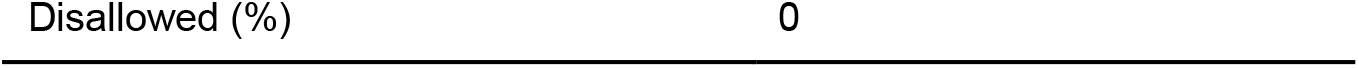
Statistics of data collection atomic refinement.

|  | **Tau-RNA fibrils (PDB:**<br>**7SP1)** |
| --- | --- |
| *Data collection and processing* |  |
| Magnification | ×130,000 |
| Voltage (kV) | 300 |
| Electron exposure (e<sup>-</sup> Å<sup>-2</sup>) | 52 |
| Defocus range (μm) | 0.76–3.9 |
| Pixel size (Å) | 1.06 |
| Symmetry imposed | C1; Helical |
| Helical Twist | 179.16 |
| Helical Rise | 2.40 |
| Initial particle images (no.) | 234,037 |
| Final particle images (no.) | 26,061 |
| Map resolution (Å) | 3.4 |
| FSC threshold | 0.143 |
| Map resolution range (Å) | 200–3.4 |
| *Refinement* |  |
| Initial model used | De Novo |
| Model resolution (Å) | 3.4 |
| FSC threshold | 0.5 |
| Model resolution range (Å) | 200–3.4 |
| Map sharpening *B* factor (Å<sup>2</sup>) | -233.0 |
| *Model composition* |  |
| Non-hydrogen atoms | 4343 |
| Protein residues | 540 |
| RNA residues | 20 |
| *B* factors (Å<sup>2</sup>) | 84.4 |
| *Protein* |  |
| *R.m.s. deviations* |  |
| Bond lengths (Å) | 0.007 |
| Bond angles (°) | 1.435 |
| *Validation* |  |
| MolProbity score | 1.75 |
| Clashscore | 11.67 |
| Poor rotamers (%) | 0 |
| *Ramachandran plot* |  |
| Favored (%) | 97.1 |
| Allowed (%) | 0 |

**Fig. 3.**
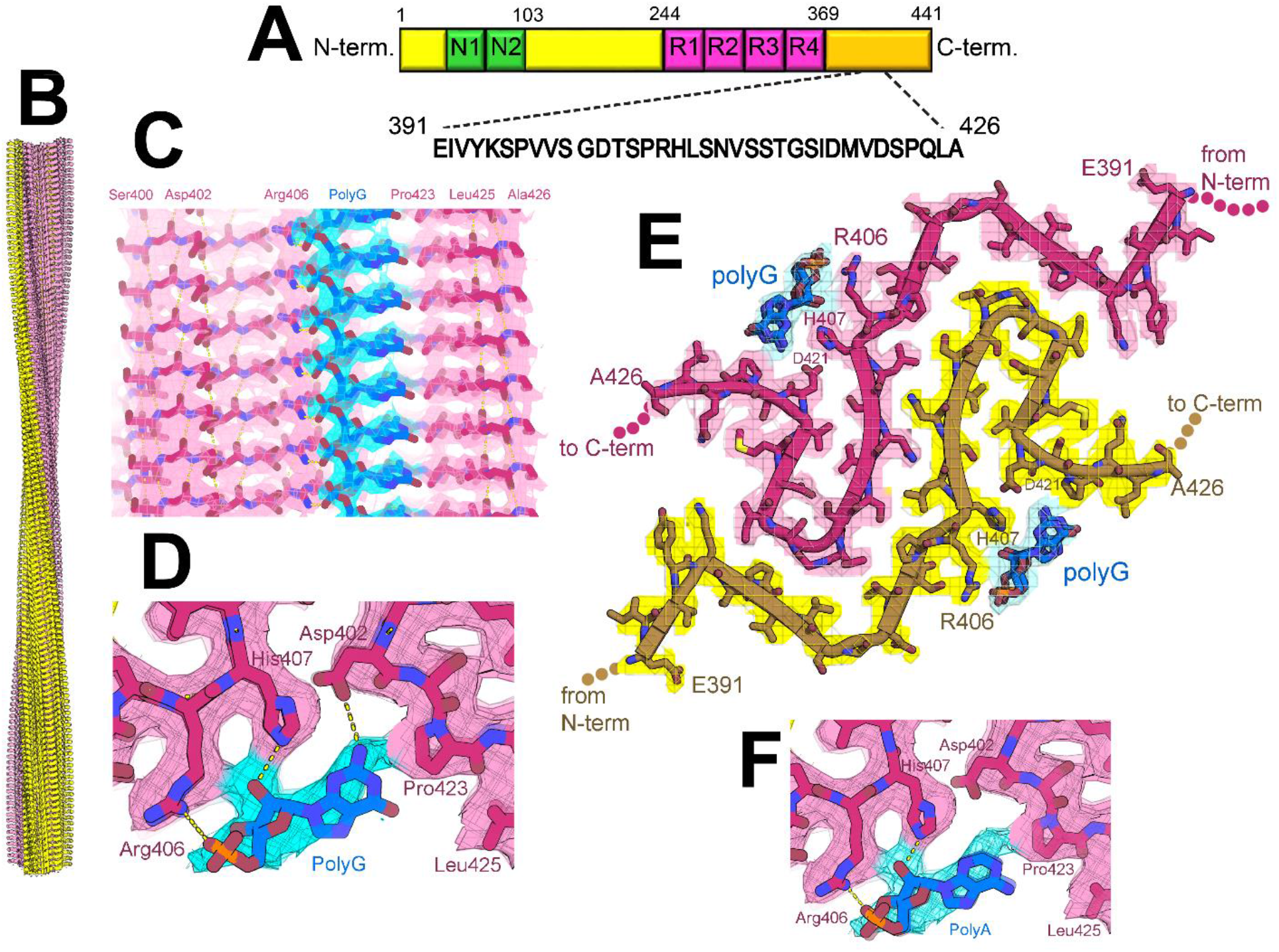
Cryo-EM structure of full-length recombinant tau fibril bound to RNA. **(A)** Schematic representation of full-length tau (tau40, residues 1-441) including the two alternatively spliced N-terminal domains and four microtubule binding domains (R1-R4). The sequence of the ordered core in our tau-RNA fibril structure is shown in black. (**B**) The CryoEM reconstruction of our tau-RNA fibril, showing two identical protofilaments (pink and yellow) with left-handed twist. **(C)** Parallel, in register alignment of tau molecules (pink) and polyG RNA (cyan) running parallel to the fibril axis. **(D)** Close-up view of tau fibril-RNA model showing H-bonding between Arg406, His407 and Asp402 and residues and polyG RNA. **(E)** Atomic model and density map of one cross-sectional layer of the tau fibril core viewed down the fibril axis. **(F)** Close-up view of tau fibril-RNA interaction showing modeled H-bonding between Arg406, His407 residues and polyA RNA.

Stripes of residual cryoEM density run along the length of the two tau protofilaments, resembling RNA strands on the fibril surface (**Fig. 3C & D**). Within each stripe, we see three adjacent blobs of density resembling a phosphate, ribose, and base moiety repeating every 4.8 Å along the fibril axis, commensurate with the repeat of stacked tau molecules. The density is not sufficiently strong to clearly distinguish among the 4 nucleotide bases, so we modeled polyA and polyG (separately) into the residual density (**Fig. 3C & F**). PolyA has been recently reported to be a cofactor for tau aggregation and localized with nuclear and cytosolic tau aggregates in P301S mouse brains (3, 29, 30).The location of our docked RNA chain is chemically compatible with the surface of the tau fibril where it is docked: (i) the phosphate OP1 atoms are positioned 2.8-3.0 Å from Arg406 NH1 atoms, suggesting they are hydrogen bonded; (ii) the ribose 2`-OH atoms are positioned 2.7 Å from His407 NE2 atoms, suggesting these atoms are also hydrogen bonded, and the adenine base contacts His407 and Pro423 (**Fig. 3C & D**); (iii) arginine-rich motifs are common among RNA binding domains including human ribosomal protein L7, the lambdoid bacteriophage N protein and HIV-1 Rev protein (37-40).

The interaction of RNA and side chains in our tau-RNA fibril is reminiscent of the interaction between side chains and unidentified cofactors in recent brain-derived fibril structures. The proximity of residual density near two positively charged side chains (Arg406 and His407) in our tau-RNA fibril matches the pattern seen in cryoEM maps of *ex-vivo* α-synuclein fibrils (41). For example, residual density is located near surface exposed Lys58 and Lys60 sidechains, and near Lys32 and Lys34 in MSA case 2 type II-1 (6xyp). Similarly, in tau fibrils extracted from CTE patients, residual density is prominent near the adjacent Lys317 and Lys321 side chains (6nwq). Hence, it is a common pattern among amyloid fibrils that adjacent pairs of positively charged sidechains cling to putative cofactors of compensating negative charge.

We note that density for RNA is relatively weak. One explanation for this weakness is the potential mismatch in distance between repeating units of tau and RNA. Tau molecules stack in a protofilament with a repeat distance of 4.8 Å, whereas the preferred base stacking distance of RNA nucleotides is 3.3 Å. As a result, the nucleotide bases stack with less aromatic overlap than normally observed in canonical A-form RNA structures. The lack of overlap probably limits the stability of the RNA-tau interaction.

### Tau-RNA fibrils are reversible amyloid

To determine whether RNA is integrated in the fibril architecture or plays only a catalytic role in amyloid fibril formation, we exposed mature fibrils to enzymatic digestion. Tau-RNA fibrils were digested by mixing the fibrils with RNase at 1: 0.6 and 1: 3 molar ratios (fibrils: RNase) for 2 hours at 37 °C. As shown in **Fig. 4A**, tau fibrils break down into small pieces with increasing RNase concentrations, suggesting that the RNA is a molecular glue for the fibril scaffold and an integral part of the fibril structure. Based on these findings, we conclude that RNA-induced tau fibrils are reversible amyloid and disaggregate when RNA is digested.

**Fig. 4.**
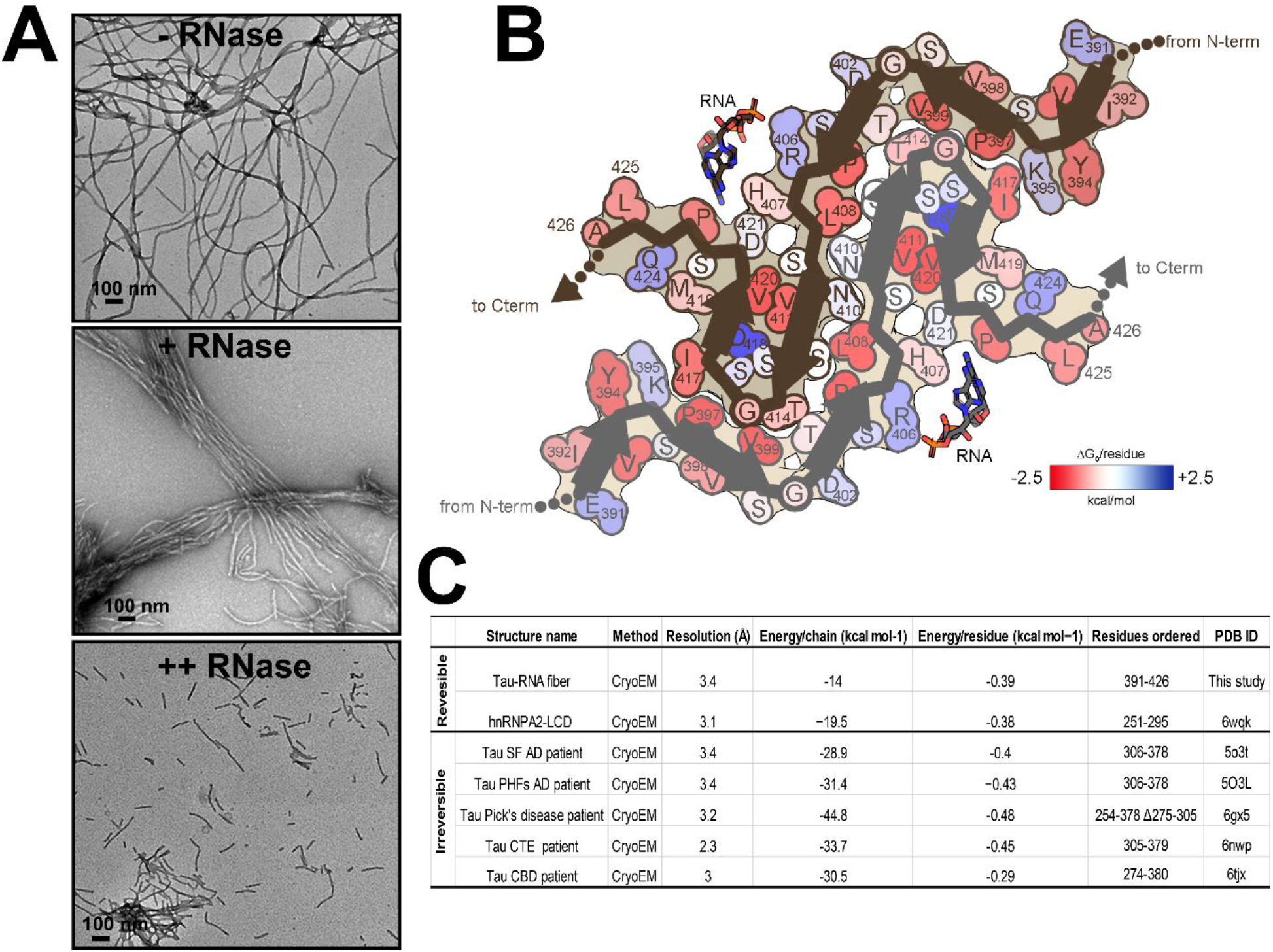
Tau-RNA fibrils are reversible amyloid. **(A)** Influence of RNase on tau fibril stability. EM image of tau-RNA fibrils in the absence of RNase (Top panel). Tau fibrils begin to cluster and breakdown in the presence of RNase at 1: 0.6 molar ratio (Fibrils: RNase, middle panel). Fibrils treated with higher molar ratio of RNase (1:3) break down into short fragments (Bottom panel). RNase was incubated with tau fibrils for 2 hours at 37 °C in 20 mM ammonium acetate buffer, pH 7.0. Scale bar 100 nm. **(B)** Solvation energy maps of tau-RNA fibrils ordered segment. Residues are colored according to their stabilization energies from unfavorable (blue, +2.5 kcal mol^−1^) to favorable (red, −2.5 kcal mol^−1^). **(C)** Comparison of the solvation energy values of the tau-RNA fiber structure with other amyloid structures.

To elucidate the structural features that impart reversibility to tau-RNA fibril assembly, we calculated a stabilization energy map of our structure (**Fig. 4B**). Our formulation of stabilization energy derives primarily from the penalty associated with dissolving and exposing atoms to water. Large negative values imply stable assemblies, difficult to dissolve. The modest stabilization energy of our fibril structure (−14 kcal mol^−1^ chain^−1^, −0.39 kcal mol^−1^ residue^−1^) is similar in value to other reversible amyloids such as hnRNPA2 low-complexity domain (−19.5 kcal mol^−1^ chain^−1^, −0.34 kcal mol^−1^ residue^−1^) and FUS (−12.2 kcal mol^−1^ chain^−1^, −0.20 kcal mol^−1^ residue^−1^) (42, 43) (**Fig. 4B & C**). FUS and hnRNPA2-LCD belong to a group of RNA-binding proteins known to phase separate and function in membraneless organelles. Pathological versions of hnRNPA2 and FUS fibrils are implicated in disease (22, 44). In contrast to reversible fibrils, more favorable stabilization energy values are observed among *ex vivo* pathological fibrils such as human serum amyloid A (−34.4 kcal mol^−1^ chain^−1^, −0.64 kcal mol^−1^ residue^−1^) (4, 45) and the brain extracted tau fibrils from patients of AD, PiD, Chronic traumatic encephalopathy (CTE) and Corticobasal degeneration (CBD) (**Fig. 4C**).

The relative ease of dissolving tau-RNA fibrils may be partly attributed to the lack of a sizable hydrophobic core. Note the paucity of strongly stabilizing residues (**Fig 4B**. bright red colors) and also the burial of an acidic residue, Asp418 (dark blue color). The interface between protofilaments is composed primarily of polar residues (Lys395, Thr403, Asn410, Ser412, Thr414); their modest contribution to stability is manifested as faint pink and blue colors in **Fig. 4B**. Most notably, Asp418 is buried and has no compensating charge nearby (**Fig. 4B**, dark blue color). Almost certainly, it is protonated in the fibril or else charge repulsion between neighboring Asp418 residues would strongly destabilize the fibril. An analogous buried glutamate (Glu8) is attributed to imparting reversibility in β-endorphin fibrils (46), and three buried aspartates in glucagon likely impart its proclivity to fibril formation at low pH (47). It seems likely that, in addition to RNA, protons may be considered to be cofactors for assembly of this fibril.

To test the capability of tau fibrils in the presence of RNA to disassemble upon raising the pH value, tau fibrils were incubated with 30 mM CAPS at pH 9.5 for 3 hrs at 37 °C. The abundance of tau fibrils in electron micrographs decreased approximately 5-fold at pH 9.5, supporting the validity of the hypothesis that pH elevation deprotonates Asp418 and Asp421 and facilitates tau fibril disassembly (**Fig. S5**).

### CryoEM structure of tau-RNA fibril reveals five steric zipper interfaces

The cryoEM reconstruction of the tau-RNA fibril reveals tight-fitting interfaces between its β-sheets that entirely exclude water molecules. Such stable structural motifs consisting of mated β-sheets are termed steric zippers. Our RNA-induced tau fibril is stabilized by five steric zippers, referred to as A_1_, A_2_, B_1_, B_2_, C (**Fig. 5A**). Steric zippers A_1_ and A_2_ are symmetrically identical zippers residing in protofilaments 1 and 2, respectively. Each is composed of a pair of β-sheets, strengthened by van der Waals contacts between side chains of one sheet face (Asp418, Val420 and Asp421) and its mated sheet face (His407, Ser409, Val411 and Ser413). Steric zippers B_1_ & B_2_ are composed of a surface displaying residues Pro397, Val399, and Thr403 mated with a complementary surface displaying residues Thr414, Gly415, and Ile417. Steric zipper C differs from the preceding zippers because it mates together identical β-sheets from separate protofilaments (homotypic). β-sheet residues Leu408, Asn410, Ser412 mate with identical residues of the opposing protofilament. In agreement with our cryoEM model, the potential for forming such a homotypic steric zipper was predicted by our ZipperDB database. Rosetta calculates energetic stabilities for homotypic hexapeptide segments. The segment 408-LSNVSS - 413 exceeds the −23 kcal/mol threshold for identifying amyloid fibril-forming segments (by 1.4 kcal/mol).

**Fig. 5.**
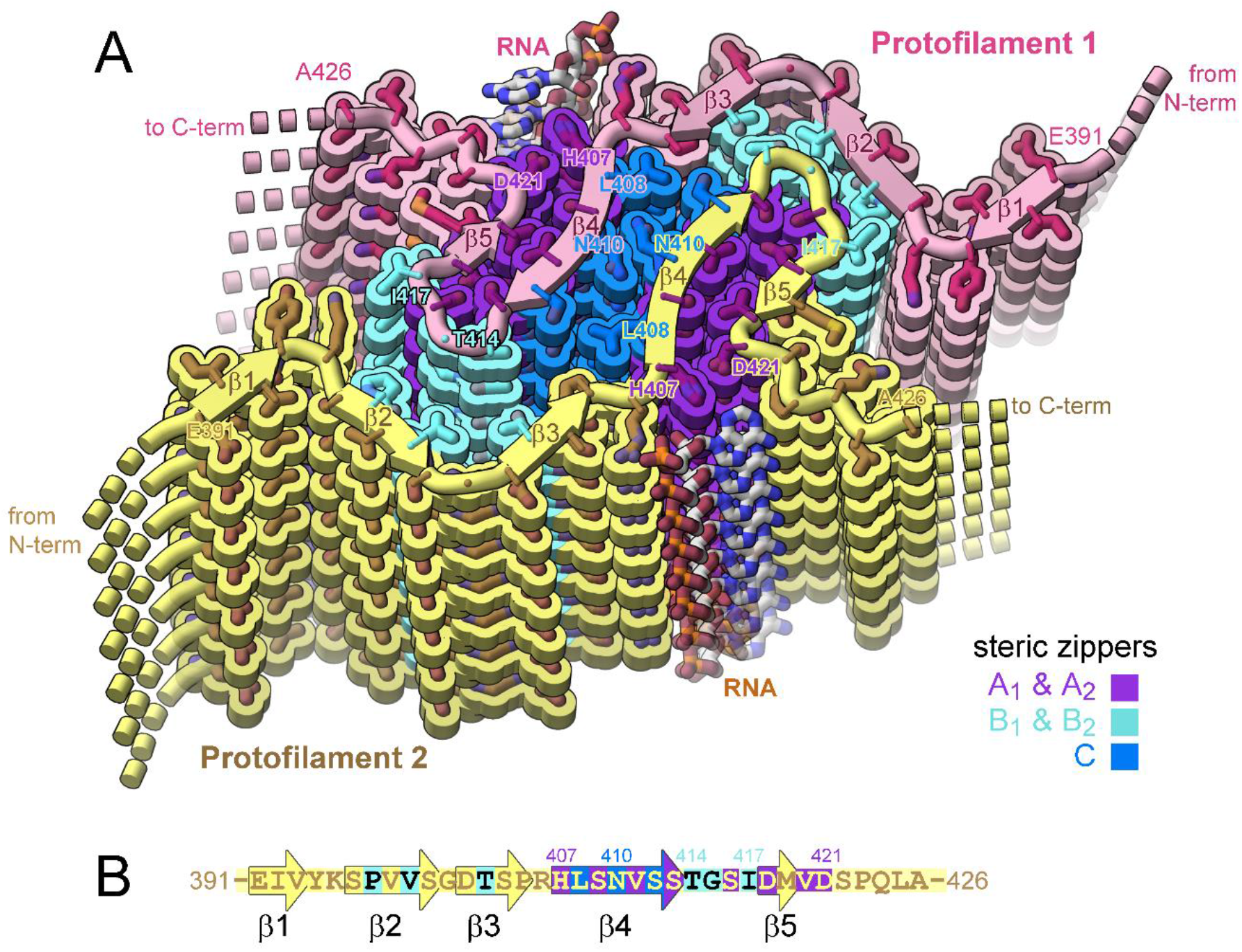
Structure of tau-RNA fibril. **(A)** Cartoon of the two identical protofilaments (pink and yellow) of tau-RNA fibrils. Eight tau layers of each protofilament are shown. β-sheet surfaces are mated together in five steric zipper interfaces (interface: A_1_, A_2_, B_1_, B_2_, and C, colored by letter). **(B)** The amino acid sequence of the ordered fibril core of tau-RNA cryoEM structure. The RNA-tau fibril core is composed of residues Glu391 to Ala426 arranged on a cross-β scaffold. The steric zipper interfaces involve β-strands 2, 3, 4, and 5 (β1: Glu391-Val393, β2: Ser396-Ser400, β3: Asp402-Pro405, β4: His407-Ser413 and β5: Asp418-Val420) that stack in layers perpendicular to the fibril axis.

Notice that the polyanion RNA interacts with tau basic residues of Arg406 and His407, thereby stabilizing the β-arch motif (residues 408-421) at the C-terminus of the ordered tau segment (**Fig. S6**). The hydrophobic interactions between the side chains of Val411 and Val420 stabilize steric zippers B1 and B2, and helps assemble the RNA binding site. The tau-RNA structured core, residues His407-Ser422 adopt a beta-arch motif, formed by multiple copies of antiparallel protein strands (β4 and β5), which stack in layers perpendicular to the fibril axis (**Fig. S6**).

### RNA stimulates conversion of tau microtubule binding repeats into fibrils

To test whether RNA may induce tau fibril formation by binding to other regions of the tau molecule than visualized in our tau-RNA structure, we tested fibrillation of three truncated versions of tau: tau-K18 (residues 244-372 with Repeat 2), tau-K18+ (244-380 with Repeat 2), and tau-K19+ (244-380 without Repeat 2) (**Fig. S7 & S8**). Tau-K18+/K19+ include an additional 8 residues C-terminal to tau-K18 in order to encompass all residues found in the core of AD fibril structures (residues 306-378). All three of these constructs lack the C-terminal domain of tau that is ordered in our full-length tau-RNA structure. EM micrographs show that addition of RNA facilitates fibril formation of 50 μM or 100 μM tau-K18 (**Fig. S7**). Interestingly, the yield of fibrils generated by tau-K18 with RNA is lower than the yield from tau40 (with RNA). The tau-K18 fibril yield increases when seeded by tau40-RNA fibrils, indicating these C-terminus-containing seeds can coax the C-terminus-deficient tau-K18 monomers into fibrils. In addition to tau-K18, we found that the constructs tau-K18+ and tau-K19+ have a greater amount of fibril formation in the presence of RNA compared to tau-K18. EM data also indicates that seeding with tau40-RNA can lead to faster and more abundant fibril formation **(Fig. S8)**. These data suggest that RNA may be able to bind to regions in the microtubule binding region of tau and promote fibril assembly. Although the fact that residues from K18, K18+, and K19+ are not found in our full-length tau-RNA structure indicates the strongest RNA-tau binding site is that visualized in our structure: Arg406 and His407.

## Discussion

Interactions of RNA with amyloid fibrils are common in both normal metabolism and pathogenesis, yet little has been uncovered about the structural basis of these complexes. To help to fill this void, we have formed amyloid fibrils of full-length recombinant tau protein with unfractionated RNA from mouse liver. The cryoEM structure of these fibrils reveals a familiar cross-beta scaffold formed by a two-protofilament tau 36–residue core. RNA runs parallel to tau’s helical fibril axis, interacting with positively charged tau sidechains (**Fig. 3C)**. Because the periodic spacing of the RNA sidechains is not commensurate with the 4.8 Å spacing of tau amyloid layers, the RNA structure is blurred, but evidently covalently intact. Its integrity is shown by the disruption of adding RNase, which causes disaggregation and dissolution of tau.

The ordered core of tau-RNA fibrils is composed of parallel, in-register β-sheets folded to form five steric zipper interfaces that stabilize the fibril, together with the extensive network of backbone hydrogen bonds (48). However, these zippers lack strong hydrophobic character. Most contacts between β-sheets involve a polar side chain **(Fig. 4B)**. Moreover, the core size is small, only 37 residues (compared to 73-107 residues in other tau fibril structures). Calculations suggest the stabilization energy of the tau core is poor (**Fig. 4C**). RNA binding appears to stabilize the tau core by contributing electrostatic and cation-π interactions with Arg406 and His407. Indeed, our fibrils degrade into short segments upon RNase treatment. Disaggregation of tau fibrils upon cofactor removal has also been observed by others (18). The polymeric connectivity of the RNA appears to be the crucial feature of its stabilization of tau fibrils because pre-digested RNA is unable to coax tau monomers into a fibril. This evidence suggests that RNA strands act as molecular glue for stabilizing the fibril **(Fig. 4A)**. All of these observations are consistent with the hypothesis that RNA can serve as a structural cofactor for reversible amyloid fibril formation of tau protein.

In addition to studies *in vitro* (6, 29, 49), other observations suggest that RNA acts as a cofactor for tau aggregation *in vivo*. RNA staining reveals that RNA associates with tau aggregates in AD and PiD (13, 50). Also, cytosolic and nuclear tau complexes with small nuclear RNAs and small nucleolar RNAs in cell culture and mouse brain models (30). And tau undergoes liquid-liquid phase separation upon binding to tRNA in neuronal cells and then transitions to a conformation resembling pathological fibrils (49).

If we assume that tau-RNA filaments akin to that of **Fig. 5** form *in vivo*, we can imagine two outcomes: either the filaments catalyze the nucleation of formation of a pathogenic amyloid polymorph, or that they protect against pathogenic forms, as envisaged from NMR studies (33). In the first outcome, our C-terminal tau filaments could facilitate the assembly of pathogenic tau filaments by epitaxial nucleation **(Fig. 6A, B, F)** by providing a 4.8 Å spacing to template growth of a commensurately spaced pathogenic fibril. Alternatively, structural order could spread from the C-terminal core to the R1-R4 regions, creating a new pathogenic core, followed by dissolution of the original core **(Fig. 6B, D, E)**. The second possible outcome is that an ordered C-terminal core such as the one we observe, can coexist with an ordered repeat region (33) which might protect the repeat region from forming pathological conformations. We imagine that our RNA-induced fibril core could be the precursor to this larger, but benign fibril core **(Fig. 6C)**.

**Fig. 6.**
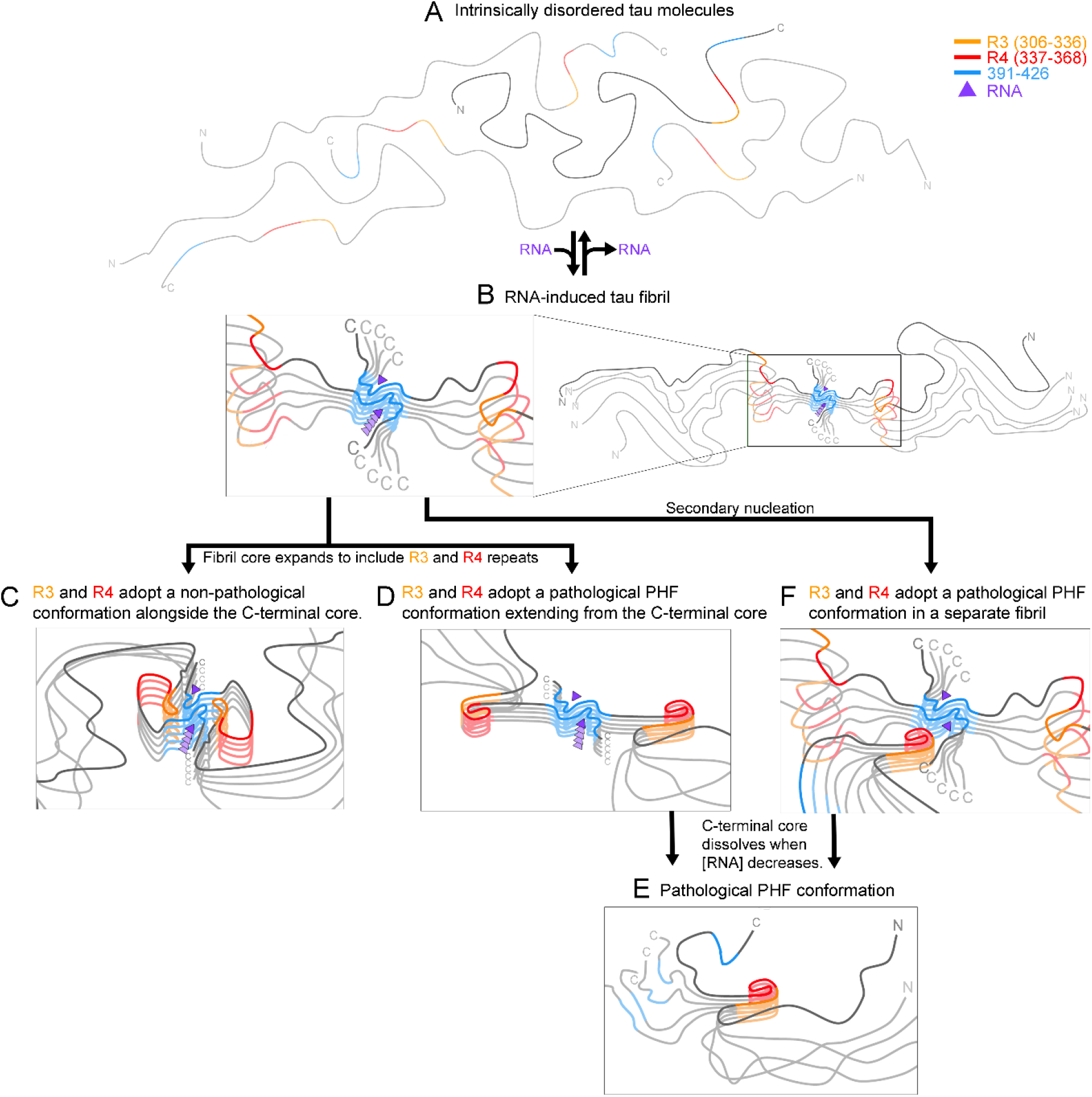
RNA-induced tau fibrils might be protective or they might facilitate formation of pathogenic tau conformations. **(A)** When intrinsically disordered tau molecules become concentrated in the presence of RNA, **(B)** RNA induces the C-terminal segment (residues 391-426) to form an amyloid core. We anticipate four possible outcomes: loss of RNA causes the fibrils to dissolve, returning tau to the soluble state (A); **(C)** the amyloid-RNA core facilitates ordering of R3 and R4 which pack aside the C-terminal core in a conformation that protects tau from forming pathological conformations as suggested by Dregni et al. (33) **(D)** the amyloid-RNA core facilitates ordering of R3 and R4 in a pathological conformation, such as the “C” shaped conformation of paired helical filaments (PHF) in Alzheimer’s disease, and then the C-terminal core disassembles, leaving **(E)** the PHF conformations; or **(F)** the RNA-amyloid core epitaxially nucleates a pathological tau conformation.

In order to test the possibilities outlined in **Figure 6**, in future work several experiments could be imagined. To test whether residues outside the tau-RNA fibril core become ordered over time as depicted in **Fig. 6C & D**, a time-resolved structural study of tau-RNA fibrils could be performed. If additional residues become ordered, CryoEM maps would reveal their structure. In order to test the possibility of epitaxial nucleation as depicted in **Fig. 6F**, the structures of fibrils seeded by tau-RNA could be studied. Indeed, our experiments already lend some support to the templating of different tau structures via the tau-RNA structure we determined here. Full-length tau, K18, K18+, and K19+ are all able to be seeded by full-length tau-RNA. Although seeding of full-length tau in the presence of RNA may result in the replication of the structure of the seed, seeding of K18, K18+, and K19+ would all result in a different fibril structure than the seed due to sequence differences (K18, K18+, and K19+ do not contain the core sequence of full-length tau-RNA fibrils). This suggests that the tau-RNA fibril may facilitate epitaxial nucleation of other tau molecules into different tau fibril structures, providing for the possibility of the mechanism illustrated in **Fig. 6F**. Future work is needed to perform time-resolved structural studies of tau-RNA fibrils and to optimize the fibrils of different tau constructs fibrillated in the presence of RNA with or without seeding to test the various possibilities we outline in **Fig. 6**.

In summary, the cryo-EM structure of tau-RNA fibrils we report here offers a near-atomic-resolution view of a structured tau C-terminus and provides a structural explanation for why tau-RNA fibrils are reversible amyloid. If it turns out that this tau-RNA structure forms *in vivo,* and is an intermediate to formation of pathogenic fibrils, our structure then offers information for the design of structure-based chemical interventions of tauopathies.

## Methods

### Recombinant protein expression and purification

Human WT tau-K18 (residues 244-372), human tau-K18+ (residues 244–380 of 4R) and human tau-K19+ (residues 244–380 of 3R tau) were expressed in a pNG2 vector in BL21-Gold E. coli. Human tau40 (residues 1-441) was expressed in pET28b vector in BL21-Gold *E. coli*. All proteins were purified as previously described (51-53).

Human tau-K18, tau-K18+ and tau-K19+ purification: BL21-Gold *E. coli* cells grown in LB to an OD600 = 0.7. Cells were induced with 0.5 mM IPTG for 3 hours at 37 °C and lysed by sonication in 20 mM MES buffer (pH 6.8) with 1 mM EDTA, 1 mM MgCl2, 1 mM DTT and HALT protease inhibitor before addition of NaCl 500 mM final concentration. Lysate was boiled for 15 minutes and the clarified by centrifugation at 32,000g for 15 minutes and dialyzed to 20 mM MES buffer (pH 6.8) with 50 mM NaCl and 5 mM DTT. Dialyzed lysate was purified on a 5 ml HighTrap SP ion exchange column and eluted over a gradient of NaCl from 50 to 600 mM. Proteins were polished on a HiLoad 16/600 Superdex 75 pg in 10 mM Tris (pH 7.6) with 100 mM NaCl and 1 mM DTT, and concentrated to ~20-60 mg/ml by ultrafiltration using a 3 kDa cutoff.

Human tau40 purification: Tau40 was expressed in pET28b with a C-terminal His-tag in BL21-Gold E. coli cells grown in TB to an OD600 = 0.8. Cells were induced with 1mM IPTG for 3 hours at 37 °C and lysed by sonication in 50 mM Tris (pH 8.0) with 500 mM NaCl, 20 mM imidazole, 1 mM beta-mercaptoethanol, and HALT protease inhibitor. Cells were lysed by sonication, clarified by centrifugation at 32,000g for 15 minutes, and passed over a 5 ml HisTrap affinity column. The column was washed with lysis buffer and eluted over a gradient of imidazole from 20 to 300 mM. Fractions containing purified tau40 were dialyzed into 50 mM MES buffer (pH 6.0) with 50 mM NaCl and 1 mM beta-mercaptoethanol and purified by cation exchange. Peak fractions were polished on a HiLoad 16/600 Superdex 200 pg in 1X PBS (pH 7.4), 1mM DTT and concentrated to ~20-60 mg/ml by ultrafiltration using a 10 kDa cutoff.

### Isolation of total RNA from mouse liver

Wild type mouse liver RNA was isolated with RNA TRIzol™ Reagent (Ambion by life technologies, Cat No: 15596018) according to manufacturer’s protocol. Briefly, 100 mg of mouse liver was homogenized in 1 ml Trizol reagent, extracted by 0.2 ml chloroform, precipitated by 0.5 ml isopropanol, and dissolved in 100 μl RNase-free H2O.

### Preparation of tau40 fibrils for cryo-EM

Recombinant tau40 was diluted at 50 μM in 20 mM ammonium acetate, pH 7.0 and incubated with 400 μg/ml of RNA. The amyloid fibril formation was examined using negative stain transmission EM after 2 days of shaking at 37 °C.

### *In vitro* ThT fluorescence assay

Recombinant tau40, tau-K18, tau-K18+ and tau-K19+ were diluted individually to 50 μM in 20 mM ammonium acetate, pH 7.0, 10 μM ThT, and mixed with 400 μg/ml of RNA. Protein was liquated to 3 replicate wells of a 384-well–plate (Thermo Scientific Nunc), and plates were incubated at 37 °C for 60 h with shaking. The fibrillation was confirmed by negative stain transmission EM.

### RNA digestions

Digestions of RNA were carried out by incubating RNA or tau-RNA seeds with RNase A (Invitrogen by Thermo Fisher Scientific, Lot: 00522943) in 20 mM ammonium acetate, pH 7.0 as shown in **Fig. 2 & Fig. S1**. The digestion of RNA was performed by incubating RNA or tau-RNA seeds with RNase at 4:1 ratio at 37°C for 6 hours.

### Effect of RNase on tau-RNA fibril stability

Tau fibrils were centrifuged at 159,000g for 1 hour and followed by washing twice with RNA free water. Fibrils was treated with RNase A at 1: 0.6 and 1: 3 molar ratio (Fibrils: RNase) in 20 mM ammonium acetate, pH 7.0 for 2 hours at 37 °C.

### Negative-stain transmission election microscopy

Negative-stain transmission EM samples were prepared by applying 4 μl of solution to 400 mesh carbon-coated formvar support films mounted on cooper grids (Ted Pella, Inc.). The grids were glow-discharged for 30 s before applying the samples. The samples were incubated on the grid for 1 min and then blotted off with a filter paper. The grids were stained with 4 μl of 2% uranyl acetate for 2 min and washed with an additional 4 μl of 2% uranyl acetate and allowed to dry for 10 min. The grids were imaged using a T12 (FEI) election microscope.

### Seeding in tau biosensor cells

HEK293 cell lines stably expressing tau-K18 was engineered by Marc Diamond’s laboratory at the University of Texas Southwestern Medical Center (54). Cells were maintained in Dulbecco’s modified Eagle’s medium (Life Technologies, Inc., catalog no. 11965092) supplemented with 10% (v/v) FBS (Life Technologies, catalog no. A3160401), 1% antibiotic-antimycotic (Life Technologies, Inc., catalog no. 15240062), and 1% Glutamax (Life Technologies, catalog no. 35050061) at 37 °C, 5% CO2 in a humidified incubator. Tau RNA-fibrils seeds was sonicated in a cup horn water bath for 5 min and then mixed with 1 volume of Lipofectamine 3000 (Life Technologies, catalog no. 11668027) prepared by diluting 1 μl of Lipofectamine in 19 μl of OptiMEM. After 20 min, 10 μl of fibrils were added to 90 μl of tau biosensor cells. The number of seeded aggregates was determined by imaging the entire well of a 96-well plate in triplicate using a Celigo image cytometer (Nexcelom) in the YFP channel. Aggregates were counted using ImageJ (55) by subtracting the background fluorescence from unseeded cells and then counting the number of peaks with fluorescence above background using the built-in particle analyzer. The number of aggregates was normalized to the confluence of each well, and dose–response plots were generated by calculating the average and S.D. values from triplicate measurements. For high-quality images, cells were photographed on a ZEISS Axio Observer D1 fluorescence microscope using the YFP fluorescence channel.

### CryoEM data collection, reconstruction, and model building

Two and half microliters of tau fibril solution were applied to Quantifoil 1.2/1.3 electron microscope grid which was glow-discharged for 4 minutes. Grids were blotted with filter paper to remove excess sample and plunge frozen into liquid ethane using a Vitrobot Mark IV (FEI). The cryo-EM dataset were collect on 300 kV Titan Krios (FEI) microscope with a Gatan K3 camera located at the S2C2 cryo-EM center. The microscope was operated with 300 kV acceleration voltage and slit width of 20 eV. Movies were acquired using super-resolution mode with a nominal physical pixel size of 1.06 Å per pixel^−1^ (0.53 Å per pixel in super-resolution movie frames) with a dose per frame of ~1.3 e- / Å^2^. Forty frames were recorded for each movie (total dose per image 52 e- / Å^2^). Automated data collection was driven by EPU automation software package. Motion correction and dose weighting was performed using Unblur and CTF estimation was performed using CTFFIND 4.1.8 (56). All fibril particles were picked manually using EMAN2 e2helixboxer.py (57). We used RELION to perform particle extraction, 2D classification, helical reconstruction, and 3D refinement (58, 59). Particles were extracted using a box size of 1024 and 686 pixels. We used 1024 pixel particles to perform the 2D classification and estimate the fibril pitch and helical parameters. We next performed 2D classifications with a box size of 686 pixels. Helical reconstruction was performed with a cylindrical reference (58).The 3D classification was performed using three classes and manually controlling the tau_fudge factor and healpix_order to separate the particles into good and bad classes. To obtain a higher resolution reconstruction, the best particles from 686-pixel box 3D classification were selected and helical “tubes” corresponding to good particles were extracted using a box size of 320 pixels. After several more rounds of 3D classification with refinement of helical twist and rise, we then used the final subset of particles to perform high-resolution gold-standard refinement. The final overall resolution estimate was evaluated to be 3.4 Å based on the 0.143 Fourier shell correlation (FSC) resolution cutoff.

### Atomic model building

The refined map was sharpened using phenix.auto_sharpen at the resolution cutoff as indicated by half-map FSC (60). We used COOT to build de novo near-atomic resolution model the sharpened map (61). The sequence registration was validated using phenix.sequence_from_map. In the final 3D refinement, we generated a five-layer model via the helical parameters and then refined the structure using phenix.real_space_refine (61). The final model was validated using phenix.comprehensive_validation (62, 63). All the statistics are summarized in **Table 1**.

### Stabilization energy calculation

The stabilization energy was calculated for each residue by the sum of the products of the area buried for each atom and the corresponding atomic solvation parameters. The overall energy was calculated by the sum of energies of all residues and different colors were assigned to each residue in the solvation energy map.

## Acknowledgments

Some of this work was performed at the Stanford-SLAC Cryo-EM Center (S2C2), which is supported by the National Institutes of Health Common Fund Transformative High-Resolution Cryo-Electron Microscopy program (U24 GM129541). The content is solely the responsibility of the authors and does not necessarily represent the official views of the National Institutes of Health.

We would to thank Dr. Hedieh Shahpasand-Kroner for mouse liver tissue at Department of Neurology, UCLA. We also thank Dr. Feng Guo at Department of Biological Chemistry, UCLA for suggestions during this work. This work was supported by Grants 1R01 AG029430 and RF1 AG054022, from the NIA National Institutes of Health. We thank Dr. Mark Arbing for preparing Tau40 bacterial pellet and purifying Tau-K19+ and the UCLA-DOE Protein Expression Technology Center which is supported by the U.S. Department of Energy, Office of Biological and Environmental Research (BER) program under Award Number DE-FC02-02ER63421.”

## Notes

### Competing Interest Statement

D.S.E. is an advisor and equity shareholder in ADRx, Inc.

