## Supplementary information for "Cryo-EM structure of RNA-induced tau fibrils reveals a small C-terminal core that may nucleate fibril formation"

### Supplementary figures

Figures S1 to S8

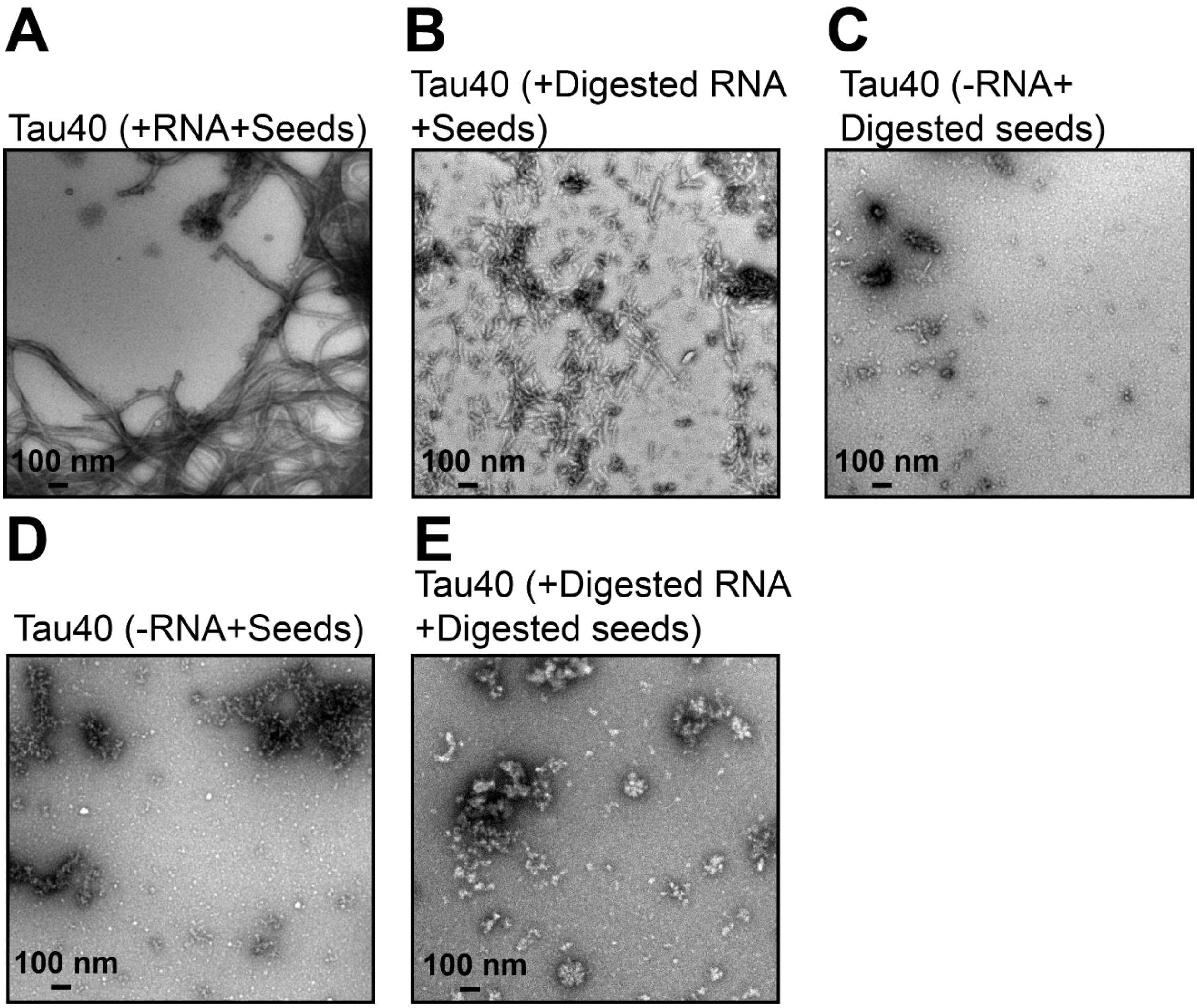


**Fig. S1. RNA seeds tau fibrils *in vitro*. (A)** Representative EM image of tau40 in the presence of seed and RNA. **(B)** EM image of tau40 in the presence of pre-digested RNA and seeds (Sonicated tau40-RNA fibrils). **(C)** EM image of tau40 in the presence of digested seeds and the absence of RNA. **(D)**The EM images of tau40 in the absence of RNA and presence of seeds. **(E)** EM image of tau40 in the presence of pre-digested RNA and digested seeds. The digestion of RNA or tau seeds was performed as described in Figure 2 legend and material and methods. Scale bar 100 nm.

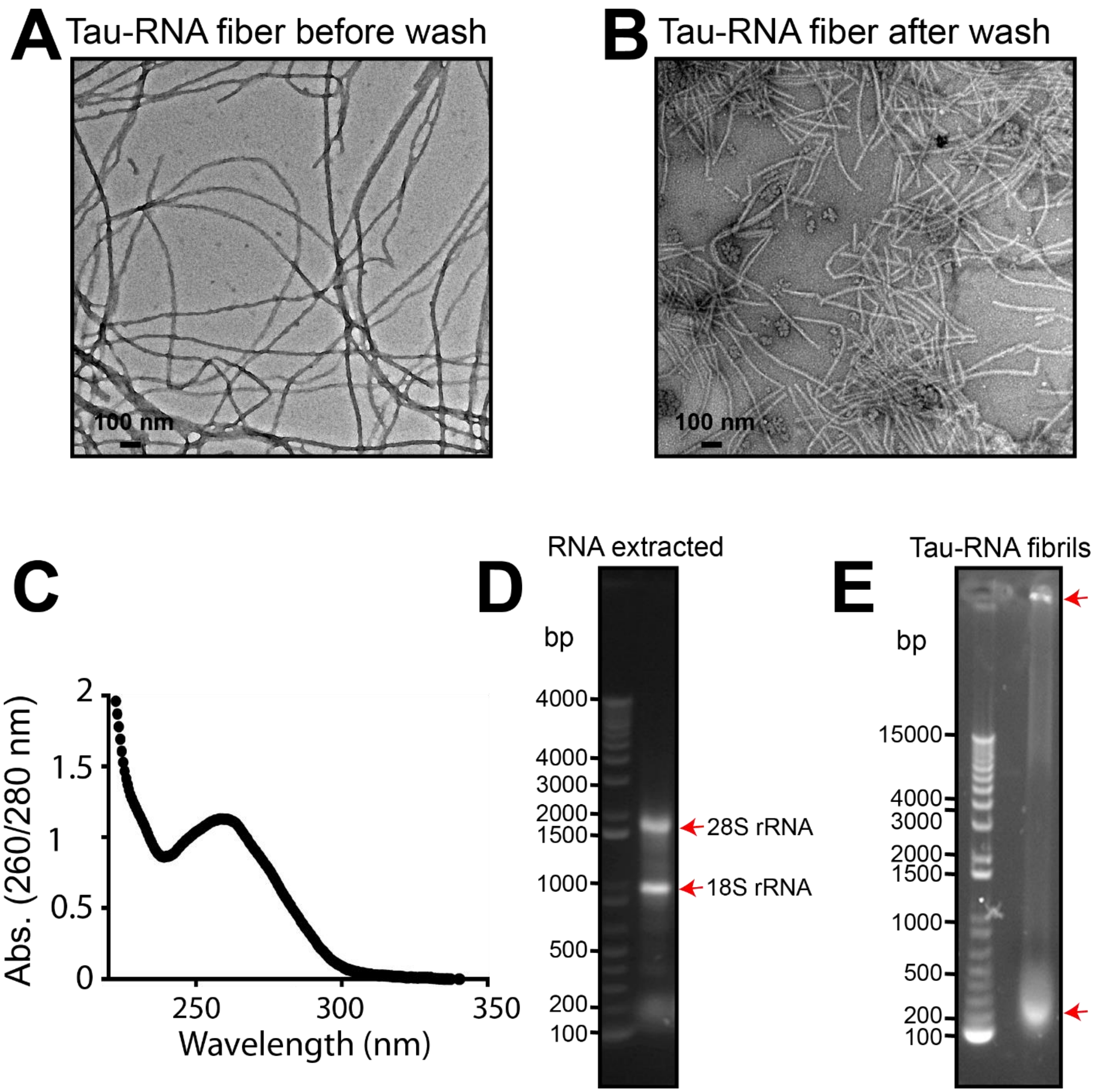


**Fig. S2. RNA is bound to tau fibrils. (A)** EM image of tau-RNA fibrils before sedimentations. **(B)** EM image of tau-RNA fibrils after sedimentation. Fibrils were washed twice with RNA-free water and centrifuged at 90,000 rpm for 1 hour. **(C)** Absorption spectra of tau40 fibril pellets shows a peak at 260 nm corresponding to RNA. **(D)** Agarose gel electrophoresis of RNA extraction. RNA extraction was performed using 1% agarose gel. We used 1 Kb Plus DNA Ladder (Invitrogen, Cat no. 10787018). **(E)** Gel electrophoresis of Tau-RNA fibrils, 5 µg of fibrils were loaded on 1% agarose. Red arrow indicates different sizes of RNA molecules associated with tau fibrils. The data indicate that the RNA is associated with tau fibrils.

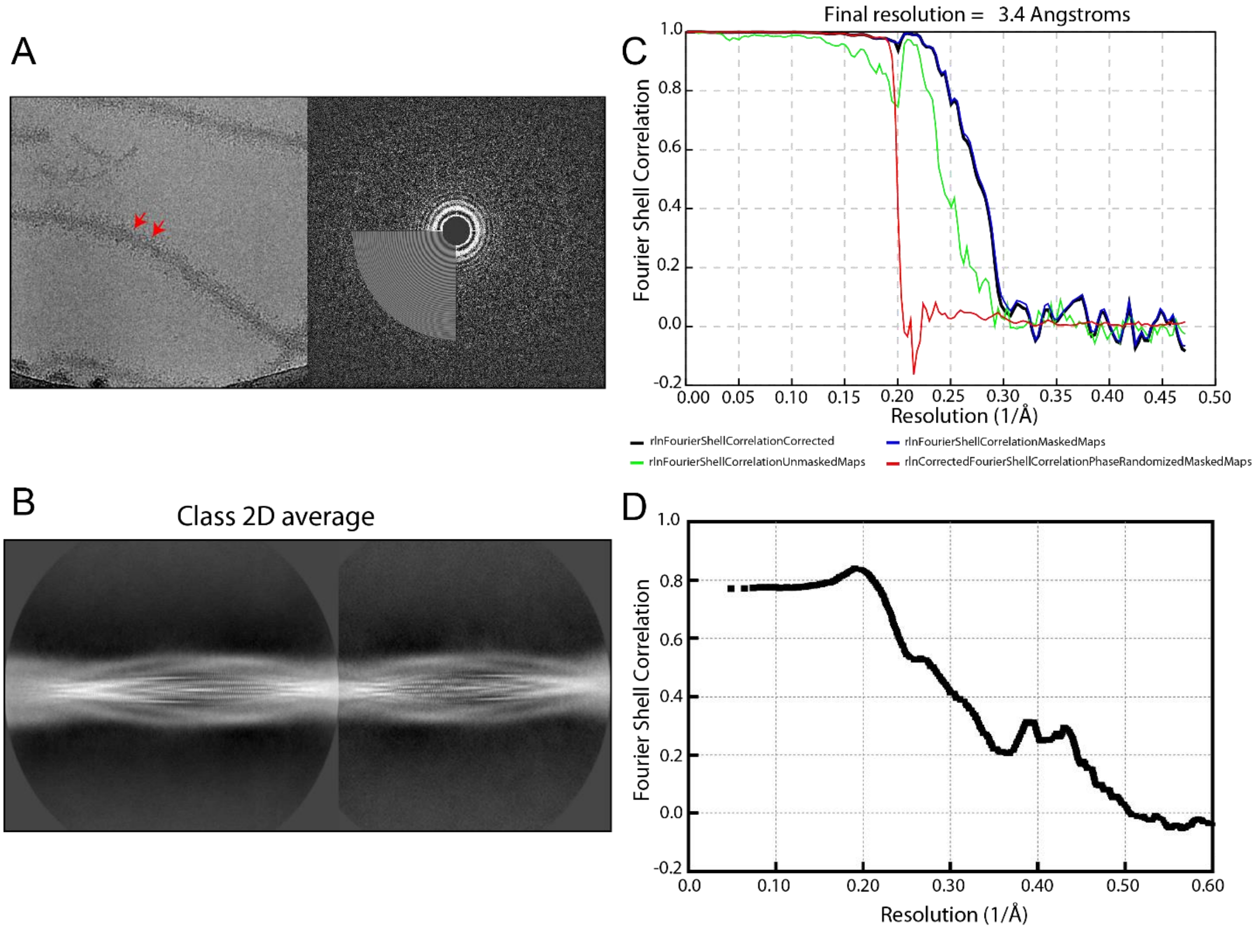


**Fig. S3.** Cryo-EM data processing. **(A)** Cryo-EM image of the full-length tau-RNA fibrils and CTFFIND diagnostic image. **(B)** 2D class averages of tau40 fibrils using 686-pixel box size. **(C-D)** FSC curves between two half-maps (C) and the cryo-EM reconstruction and refined atomic model (D).

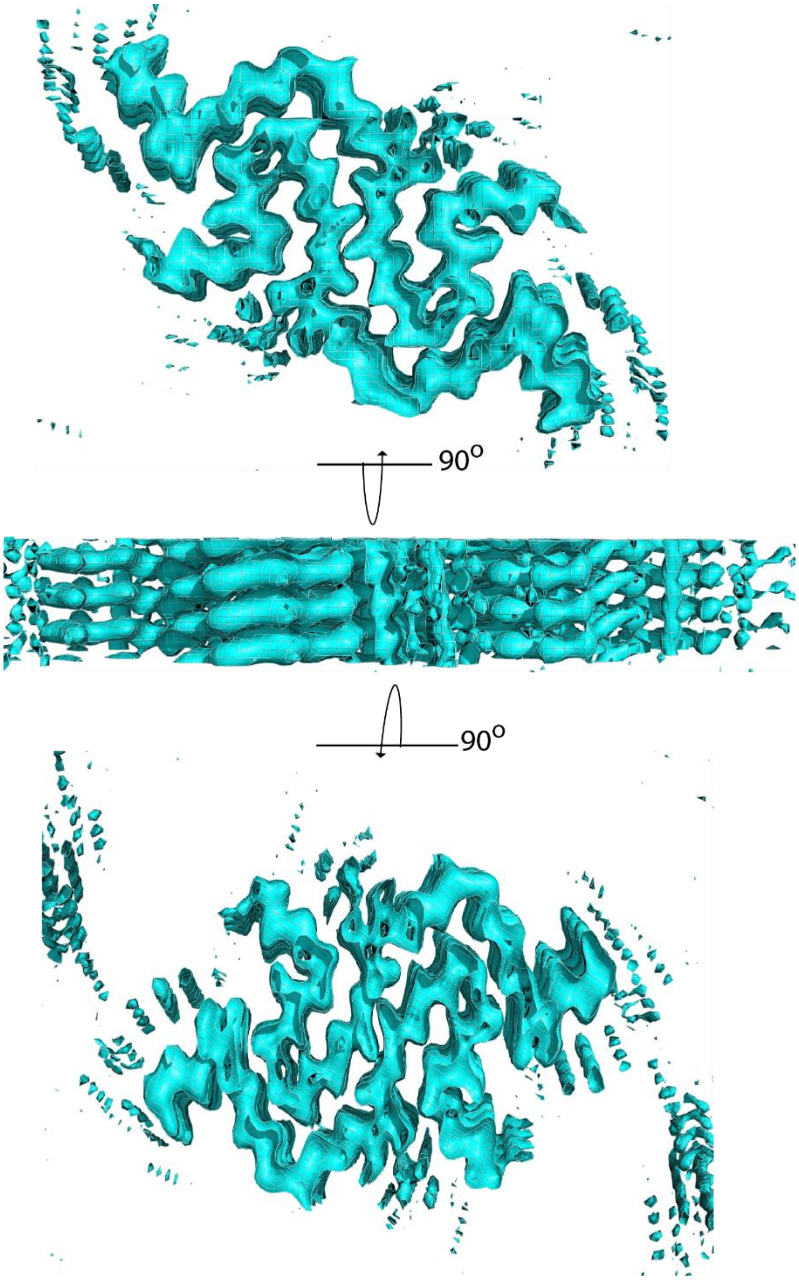


**Fig. S4.** Orthogonal views of tau-RNA cryo-EM map, covering three layers perpendicular to the fibril axis. The top and bottom panels show clear separation between β-strands (and protofilaments), and the middle panel shows clear separation between β-strands along the fibril axis.

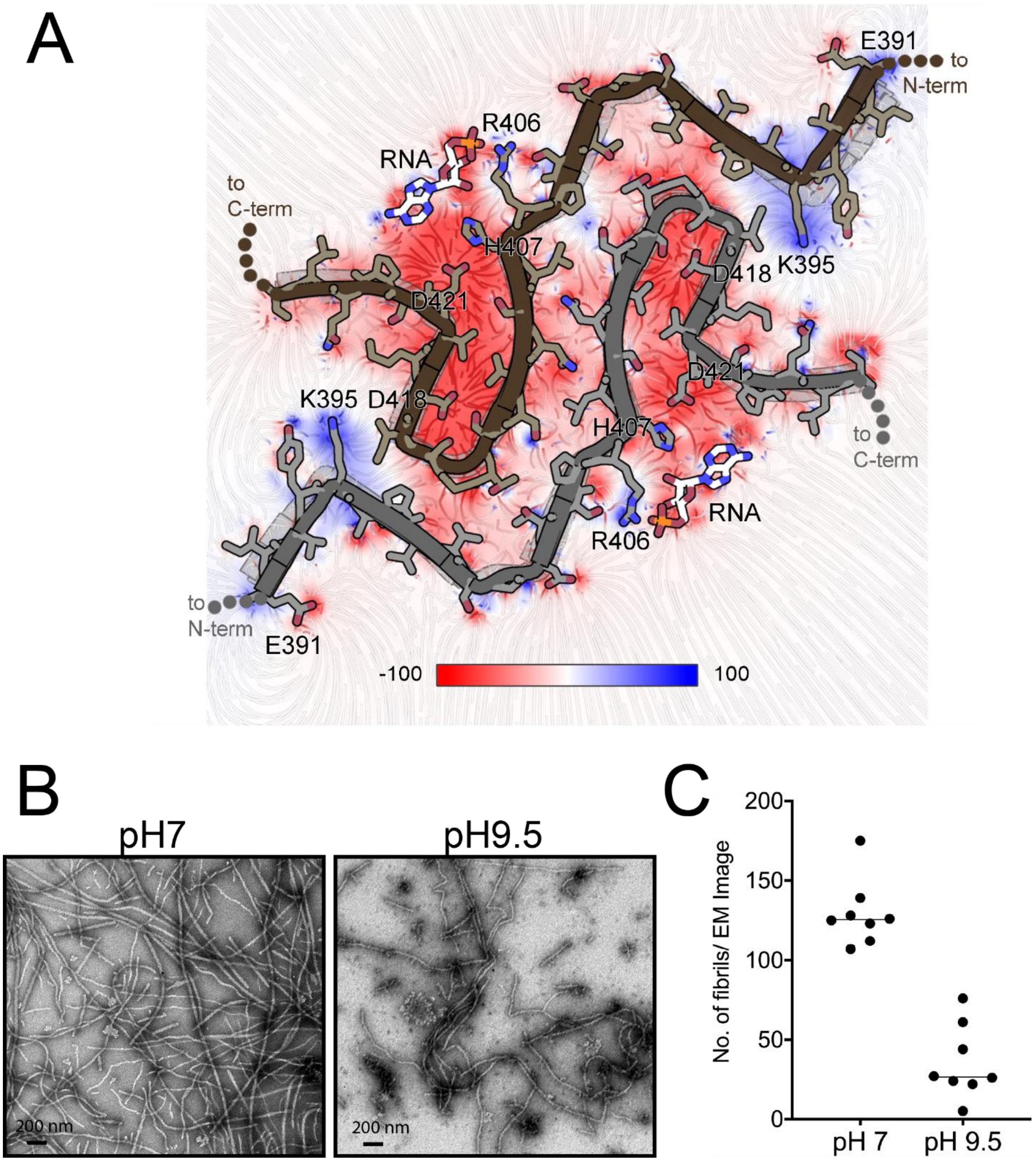


**Fig. S5. Tau-RNA fibrils show signs of disassembling at elevated pH**. **(**A) Electrostatic potential surface of tau fibrils with bound RNA. Regions of negative potential are colored red, those of positive potential are colored blue, neutral regions are shown in white. Asp418 and Asp 421 residues are likely protonated in the fibril structure, but here the electrostatics calculation program, APBS, considers them charged. Consequently, this image depicts the repulsive force that would develop at alkaline pH, and may assist fibril disassembly. Units on the scale bar are kT/e$^-$ at 300 K. **(B)** Influence of pH value on the tau fibril stability. Fibrils incubated at 20 mM ammonium acetate pH 7 at 37 ˚C for 3 hours (Left panel). The amount of tau fibrils diminished upon incubation with 30 mM CAPS pH 9.5 at 37 ˚C for 3 hours (right panel). The abundance of fibrils in these two images are representative of their respective EM grids. **(C)** Quantification of tau-RNA fibrils appearing in eight EM images collected at each of the two pH values. Each dot represents the fibril count in one EM image.

.

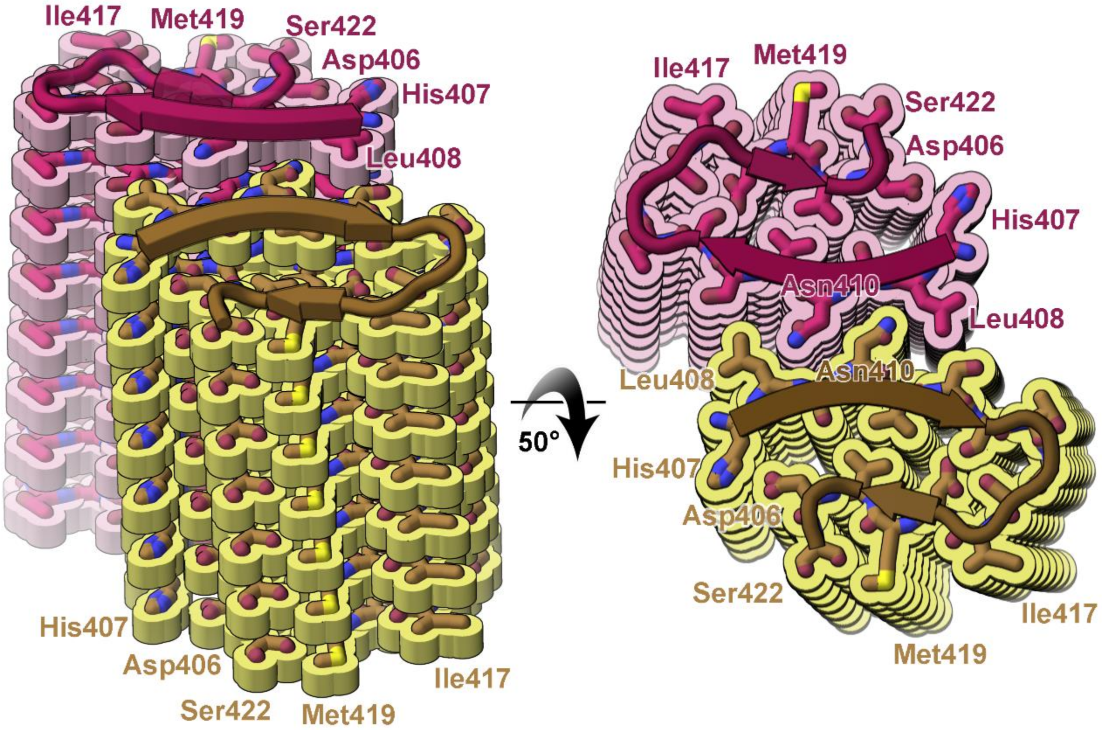


**Fig. S6. The tau-RNA fibril core is centered on pair of beta-arches.** (A-B) Ribbon representation of the beta-arch motifs. A beta-arch motif is a strand-turn-strand unit stabilized by the interaction of the side chains of the adjacent beta sheets. (A) Side view of the beta-arches. (B) Top view of the beta-arches.

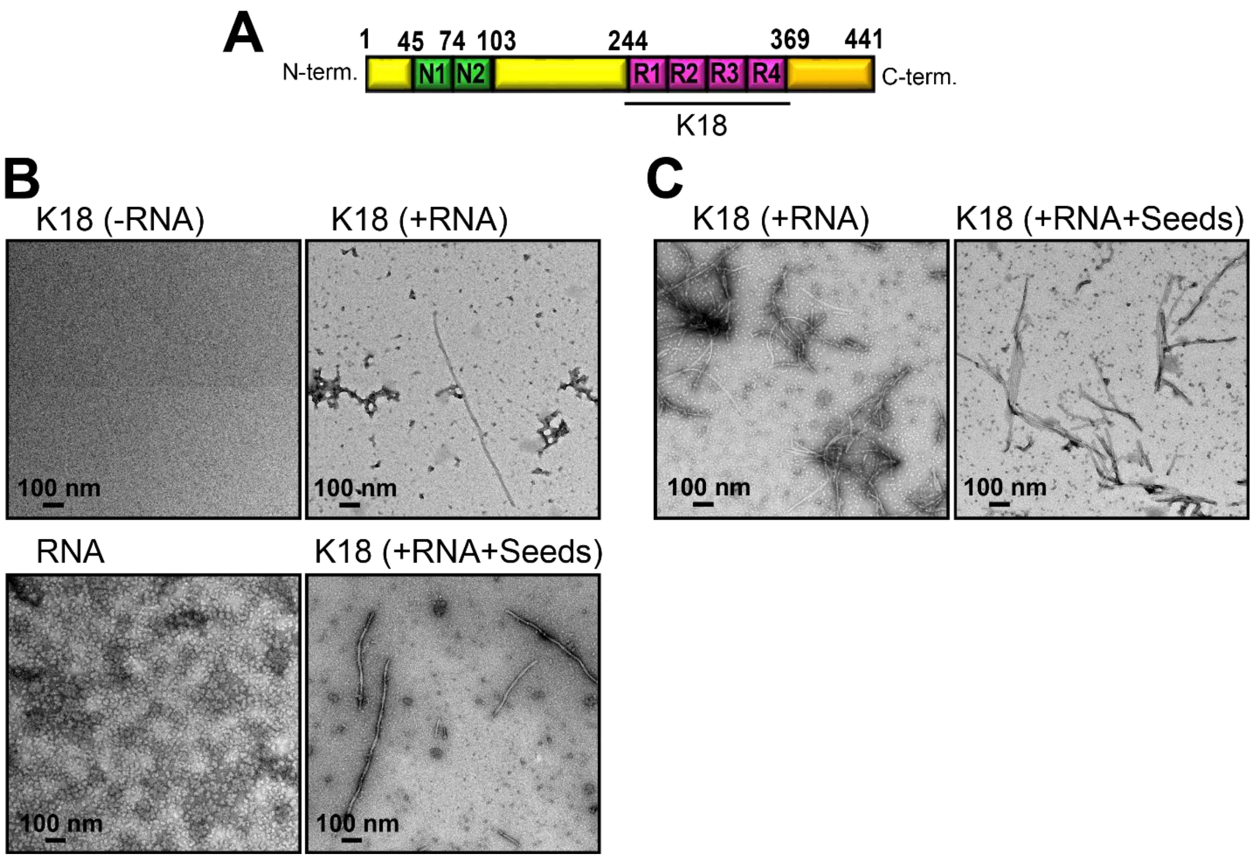


**Fig. S7. *In vitro* aggregation of tau-K18 fibrils requires RNA cofactor. (A)** Schematic representation of full-length tau (tau40, residue 1-441) including the N-terminus, C-terminus and four microtubule binding domains (R1-R4), named tau-K18 (residues 244-369). **(B)** Representative EM image of tau-K18 (50 µM) in the absence of RNA (left panel), in the presence of RNA (right panel), tau-K18 in the presence of RNA and tau40-RNA fibril seeds (right panel bottom) and EM image for RNA only (left panel bottom). **(E)** EM image of tau-K18 (100 µM) in the presence of RNA (left panel) and in the presence of RNA and seeds (right panel). Scale bar 100 nm.

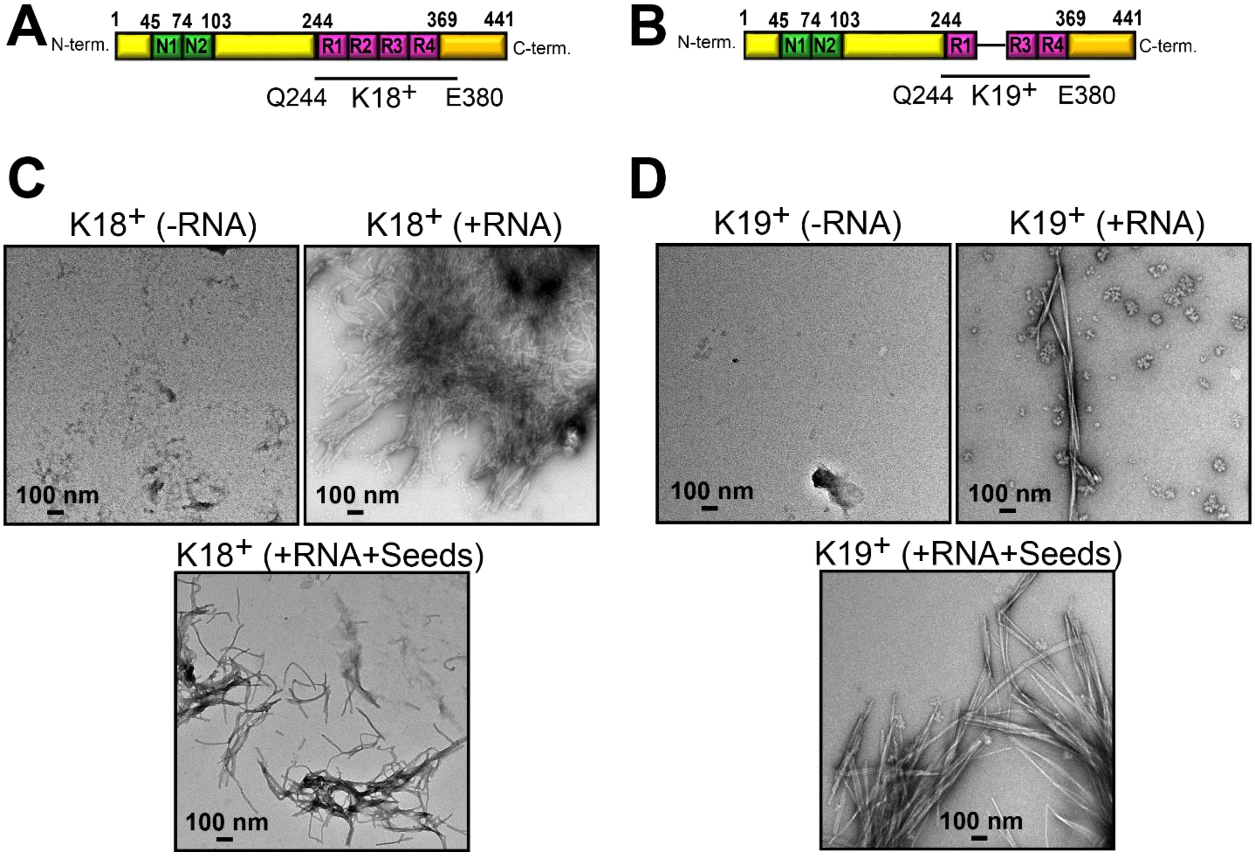


**Fig. S8.** ***In vitro*** **aggregation of tau-k18+ and tau-k19+ fibrils requires RNA cofactor.** **(A)** Schematic representation of tau-K18+ including the four microtubule binding domains (R1-R4), (residues 244-380). **(B)** Schematic representation of tau-K19+ (residues 244-380 of 3R). **(C)** Representative EM images of tau-K18+ in the absence of RNA (left panel), in the presence of RNA (right panel), tau-K18 in the presence of RNA and preformed seeds (middle panel bottom). **(D)** EM image of tau-K19+ in the absence of RNA (left panel), tau-K19+ in the presence of RNA (right panel) and in the presence of RNA and seeds (middle panel bottom). Scale bar 100 nm.
